# Memoization on Shared Subtrees Accelerates Computations on Genealogical Forests

**DOI:** 10.1101/2024.05.23.595533

**Authors:** Lukas Hübner, Alexandros Stamatakis

## Abstract

The field of population genetics attempts to advance our understanding of evolutionary processes. It has applications, for example, in medical research, wildlife conservation, and – in conjunction with recent advances in ancient DNA sequencing technology – studying human migration patterns over the past few thousand years. The basic toolbox of population genetics includes genealogical tress, which describe the shared evolutionary history among individuals of the same species. They are calculated on the basis of genetic variations. However, in recombining organisms, a single tree is insufficient to describe the evolutionary history of the whole genome. Instead, a collection of correlated trees can be used, where each describes the evolutionary history of a consecutive region of the genome. The current corresponding state of-the-art data structure, tree sequences, compresses these genealogical trees via edit operations when moving from one tree to the next along the genome instead of storing the full, often redundant, description for each tree. We propose a new data structure, genealogical forests, which compresses the set of genealogical trees into a DAG. In this DAG identical subtrees that are shared across the input trees are encoded only once, thereby allowing for straight-forward memoization of intermediate results. Additionally, we provide a C++ implementation of our proposed data structure, called `gfkit`, which is 2.1 to 11.2 (median 4.0) times faster than the state-of-the-art tool on empirical and simulated datasets at computing important population genetics statistics such as the Allele Frequency Spectrum, Patterson’s *f*, the Fixation Index, Tajima’s *D*, pairwise Lowest Common Ancestors, and others. On Lowest Common Ancestor queries with more than two samples as input, `gfkit` scales asymptotically better than the state-of-the-art, and is thus up to 990 times faster. In conclusion, our proposed data structure compresses genealogical trees by storing shared subtrees only once, thereby enabling straight-forward memoization of intermediate results, yielding a substantial runtime reduction and a potentially more intuitive data representation over the state-of-the-art. Our improvements will boost the development of novel analyses and models in the field of population genetics and increases scalability to ever-growing genomic datasets.

**2012 ACM Subject Classification:** Applied computing → Computational genomics; Applied computing → Molecular sequence analysis; Applied computing → Bioinformatics; Applied computing → Population genetics

## 1 Introduction

Charles Darwin famously wrote that living beings share a common evolutionary history and organize in a tree [6, 14]. Since then, hand-drawn trees based on morphological features [14, 20, 22] have been replaced by trees computed using mathematical and statistical models that take into account variations among the genomic data of the observed species [43, 21].

Next to analyzing the evolutionary history among distinct species (*phylogeny*), the question of how living and past individuals and populations of a single species, for example humans^1^, have emerged, migrated, and how they relate to each other (*genealogy*) is also of interest. Population genetics is the sub-field of evolutionary biology addressing these questions and advances our understanding of evolutionary processes such as mutations, genetic drift, gene flow, and natural selection [34]. Its downstream applications include for example medical research [55, 40, 32, 3], wildlife conservation [23, 34, 51, 47], and – in conjunction with recent advances in ancient DNA sequencing technology [41, 37, 45, 12] – studying human migration patterns in the past few thousand years [2, 33, 46, 41].

### 1.1 Genealogical Trees, Tree Sequences, and Forests

*Genealogical tress* describe the shared evolutionary history among individuals and are calculated on the basis of the genetic variation across sampled individuals [30]. Genetic loci (positions) where all sampled genomes are identical do not carry evolutionary signal and are therefore omitted. Hence, *genomic sequence* or *genome* in this work often refers to only those loci of the genome where there exists variation across the samples. In genealogical trees, the *tips* (out-degree 0 nodes) represent biological samples, that is, the genetic material of a single individual. We call non-tip nodes *inner nodes*; these represent one or multiple variations in the sequence inherited from a single parent. These inner nodes might be multifurcating, that is, they can have more than two children. Further, we consider all genealogical trees to be rooted, that is they have a node with an in-degree of 0 (i.e., no parent), which we call the *root*, and directed edges going away from this root. The root of a genealogical tree is the hypothetical common ancestor individual of all tips in the tree.

Single trees cannot adequately describe processes which cause individuals to pass on different parts of the genome independently [38]. This is the case, for example, in sexually reproducing species [24]: Here, a process called *recombination* causes an offspring to inherit parts of its genome from its biological father and other parts from its biological mother. During recombination, the genetic material, organized in a DNA strand, breaks and reconnects. Thus, the closer two genetic positions are along the genome, the higher the chance that they have both been inherited from the same parent [50]. Therefore, for recombining organisms such as humans, inferring and using different (correlated) trees to model the (correlated) history of distinct regions along the genome captures their evolutionary history more accurately than a single tree [24]. Based on this observation, Kelleher et al. [29, 30, 44] introduce so-called *tree sequences* (Figure 1.a), which are collections of genealogical trees, each describing the evolutionary history of a range of adjacent sites along the genome. Conceptually, tree sequences are closely related to Ancestral Recombination Graphs [57]. In the genealogical trees stored in a tree sequence, tips represent biological samples, edges represent lines of descent, and mutations are meta-data stored at these edges (Figure 1). In tree sequences, the edges are valid for a specified region of the genome, as they describe its evolutionary history. Statistics over the topologies of genealogical trees (Section 1.2) and differences in the genetic code between the observed individuals (Section 1.3) are frequently deployed in population genetics to conduct quantitative assessments, e.g. how genetically diverse a population is.

**Figure 1.**
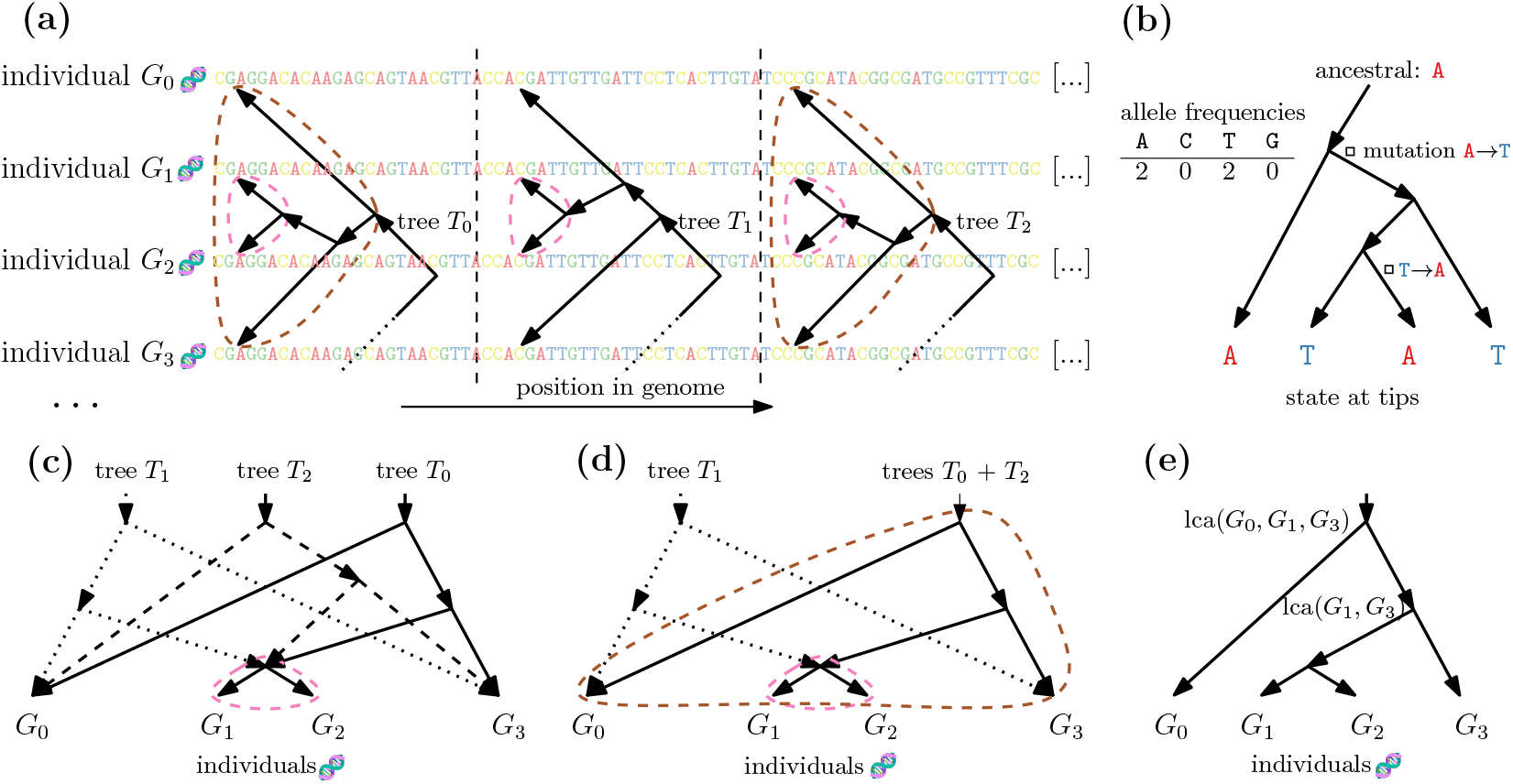
**(a)** Different, but correlated, trees each describe distinct regions of multiple aligned genomes. The subtree (*G*_1_,*G*_2_) (pink) appears in all displayed trees, whereas the subtree (*G*_0_,((*G*_1_,*G*_2_),*G*_3_)) (bronze) appears only in *T*_0_ and *T*_2_, but not in *T*_1_. **(b)** A tree describes the genomic states of genomes at its tips by storing ancestral states and corresponding mutations annotated at the tree edges (we show only a single genomic site). **(c)** `tskit` encoding of the three trees from (a). `tskit` re-uses (*G*_1_,*G*_2_) but not (*G*_0_,((*G*_1_,*G*_2_),*G*_3_)), as the latter does not re-occur in *consecutive* trees. **(d)** Our proposed encoding (`gfkit`) of the trees from (a). `gfkit` re-uses both, (*G*_0_,*G*_1_) when describing *T*_1_, and (*G*_0_,((*G*_1_,*G*_2_),*G*_3_)) when describing *T*_2_. **(e)** Lowest Common Ancestors of two (lca(*G*_1_, *G*_3_)) or more (lca(*G*_0_, *G*_1_, *G*_3_)) selected samples.

### 1.2 Lowest Common Ancestor (LCA) Queries

The Lowest Common Ancestor^2^ (LCA) [1] of two or more tips in a tree is the lowest (farthest from the root) inner node located on *all* of the paths from the tree’s root to the given tips (Figure 1.e). A population geneticist could, for example, use LCA queries on the set/sequence of genealogical trees to answer the following question: “Is the evolutionary history of individuals of group *A* and *B* separate at the same time as this land bridge vanished?”. For this, one would first compute the LCA of all individuals in *A* and *B* – and then estimate the time of the resulting ancestor using methods beyond the scope of this paper. Note that here, we extend the definition of LCA to also include more than two tips.

### 1.3 Differences in The Genetic Code Among Individuals

*Alleles* are a concrete variant of a genetic variation between individuals and encompass one or multiple (related) genetic loci. In this work, we consider only alleles consisting of genetic differences at a single genomic location. For example, a set of genomic sequences of human individuals might have the alleles `A` and `C` at a specific genome site. We call a site with more than one allele *polyallelic* (e.g., `A`, `C`, or `T`), or *biallelic* if it has exactly two alleles. The proportion of alleles at a given site is called the *allele frequency*, e.g., 80 `A`s and 5 `C`s (Figure 1.b). We are often interested in selecting only a subset of the samples (tips in the genealogical tree) for a given query; we call these the *selected* samples or the *sample set*. The exponential number of possible sample sets yields pre-computing all statistics of interest infeasible, and therefore requires algorithmic improvements.

Many statistics used in population genetics are a function of the allele frequencies [44]: For example the sequence *diversity* [39] describes the average genetic difference among the genomes of individual samples in a single sample set (drawn from a single population). Other statistics are concerned with the differences between two or more sample sets, for example to assess the degree of sequence *divergence* [39] between them. While these statistics are defined per site, we are often interested in the average value across many or all sites in the input genome or within a sliding window over all sites of the genome.

### 1.4 The Need for Efficient Algorithms

Storing and processing the full genomes of all samples contained in contemporary datasets on a base-by-base basis is infeasible. To date, there exist collections of hundreds of thousands of modern and ancient genomes for humans alone [5, 35, 11, 53, 28, 46], and even larger datasets are currently being sequenced and assembled.^3^ In fact, the amount of available genomic data is growing faster than the amount of computational power and storage per unit of money [8]. We can therefore not rely on increasing (sequential or parallel) processing speeds to analyze these data. Instead, we require algorithms which perform no more work than minimally required by the problem^4^ (i.e., are compute-efficient) as well as highly optimized implementations which exploit modern hardware features and leverage data locality.

### 1.5 State of the Art and Contribution

Tree sequences allow for reducing the storage space required for a set of related genomes by modelling the evolutionary history of the included samples, and reusing computations in statistical queries [29, 30, 44]. For example, the human chromosome 20 in the Thousand Genomes Project collection (Section 5) contains 5008 sequences with 860 thousand variant sites; each sequence being in one of four potential states (`A`, `C`, `T`, or `G`) at each site [30]. This results in a theoretical space requirement of 1 GiB if stored on a base-by-base basis using 2 bit per sequence and site – compared to the tree sequence file size of 283 MiB [30].

Additionally, a tree sequence stores the evolutionary history of each part of the genome via the corresponding genealogical trees and allows reusing partial results between adjacent, yet topologically distinct trees, thereby avoiding some – yet not all possible – redundant computations [29, 44]. This is because tree sequences reuse only intermediate results for shared subtrees among *adjacent* trees along the genome [29, 44]. They do not allow reusing intermediate results across trees further apart, if the respective subtree is not part of all intermediate trees (Figure 1.c). Tree sequences use edit operations (edge insertions and removals) to describe changes/differences in the tree topology from one tree to the next along the genome; and thus across recombination events. Therefore, determining if a given subtree was already present in an earlier tree is not trivial. For example, if an edge from node *a* to node *b* (*a → b*) is removed and a (sans meta-data) identical edge *a → b* is added back a few trees further down the genome, other edges might have changed in the subtree below node *b*, thus invalidating the intermediate results for *b*.

The core concept of our data structure is to encode and store shared subtree across *all* (not just adjacent) trees exactly once, enabling queries on the trees topologies (Section 3.3) and genetic sequences (Section 3.4) to reuse intermediate results across *all* trees (*memoization*).

### 1.6 Overview of the Paper Structure

After providing an overview of related work (Section 2), we describe the core idea of our proposed data structure `gfkit` (Section 3), as well as the implications of our design decisions. Further, we describe `gfkit`’s query algorithm for computing the LCA (Section 3.3) and other common statistics used in population genetics (Section 3.4). We also describe the conversion of the `tskit` to the `gfkit` data structure (Section 3.6) as well as two variants of our data structure (Section 3.5 and Section 3.7). Further, we evaluate the performance of `gfkit` (Section 6), report speedups (Section 6.1 and Section 6.2) and analyze the created DAGs (Section 6.3 and Section 6.4). Next, we discuss the space usage of `gfkit` vs `tskit` (Section 6.6) and describe our qualitative observations concerning numerical stability (Section 7). Finally, we conclude, and highlight possible direction of further work (Section 8).

## 2 Related Work

The genetic code of related species or individuals of the same species is highly redundant, even when considering only variant sites [44]. Ané and Sanderson [4] thus propose compressing related genomic sequences using a phylogenetic tree. Here, the tips of the tree represent the genomic sequences, which are fully described by storing the ancestral state for each site at the root and the respective mutations along the edges of the tree (Figure 1.b). Ané and Sanderson primarily use a biologically reasonable evolutionary tree with an optimized parsimony score in order to attain a good compression ratio but not for explicitly modelling evolutionary history. Kelleher et al. [29, 30, 44] introduce so-called *tree sequences*, which model the evolutionary history of a set of related genomes, allow reusing some intermediate results when computing statistics on them, and enable space-efficient storage of these sequences (Section 1.5).

Matthews and Williams [36] compress a collection of trees by encoding each *bipartition* exactly once. Here, a bipartition is a separation of a tree’s tips into two disjoint sets. Note, that each edge of a tree induces a bipartition and thus the set of all bipartitions fully describes a tree. However, this does not allow for direct access to the trees’ topologies, which we require, for example, to compute the Lowest Common Ancestor (Section 3.3).

Directed Acyclic Graphs (DAGs) contain only directed edges and no path of length ⪈ 0 from a node to itself. The tree terminology introduced in Section 1.1 generalizes to DAGs: We call out-degree 0 nodes *tips*, which represent samples and their associated genomes. Contrary to trees, DAGs may have multiple *root* nodes, that is, nodes with an in-degree of 0.

Theoretical computer science has studied the compression of single trees via DAG-compression [48, 10], tree grammars [16], and top-trees [9]. DAG-based compression reuses identical (topology and label) subtrees, when occurring multiple times in the same tree. In a tree, each node also induces/represents the subtree containing it and all of its descendants. Thus, instead of encoding a subtree a second time, we add an edge to the node representing the already encoded subtree, resulting in a DAG. In a single genealogical tree, all tips (representing samples) are distinct and thus not compressible using DAG-based compression. However, one can extend the idea of representing each unique subtree only once to forests by reusing subtrees that are part of multiple trees (Section 3). Ingels [26, 25] implements this idea, encoding a forest as a DAG. However, they focus on the enumeration of trees instead of on reusing intermediate results during computations. Tree grammars and top-tree based compression reuse tree patterns, that is any identical connected subgraph (not necessarily including all descendants) [9]. It remains an open question if top-trees^5^ can be used to encode the set of related genealogical trees while supporting the required queries efficiently.

To the best of our knowledge, two phylogenetic tools use DAGs to memoize intermediate results: ASTRAL-III, a tool for inferring a species tree from a set of topologically distinct gene trees, represents and processes unique tip bipartitions only once [59]. Further, Larget [31] uses memoization to compute the conditional clade probability only once per unique subtree.

## 3 Design of the Genealogical Forest Data Structure

In recombining organisms, the collection of genealogical trees used to describe the related evolutionary histories of different parts of the genome share common subtrees (a node including all of its descendants).^6^ Thus, we propose a data structure which encodes each unique subtree across *all* trees exactly once (Figure 1.d). We construct such a data structure by contracting the set of unconnected trees to a Directed Acyclic Graph (DAG) where each root node represents a tree and each non-root node represents a unique subtree in the input set of trees. Here, we consider each sample to be a subtree containing only itself. If two or more input trees share an identical subtree, the associated node in the DAG has multiple incoming edges; see for example `(B,C)` in Figure 1.d. These resulting DAGs are called *multitrees* [18]. In them, each subgraph and all nodes reachable from it induce a tree. Additionally, there is exactly one path from each root to each tip. In analogy to tree sequences (implemented in `tskit`), we call this data structure a *genealogical forest* and provide an implementation called `gfkit`. We choose this naming to highlight that the trees describe genealogies and use the established term “forest” to describe a collection of trees.

We encode the (closely related) genomic sequences represented by the tips of the genealogical forest analogously to `tskit` (Figure 1.b). For this, we make the biologically realistic assumption that the evolutionary history of each genomic position is described by exactly one tree. Several adjacent positions along the genome can and will often share the same tree. This assumption can potentially be relaxed - see future work (Section 8). For each site with genetic variation^7^, we store the ancestral state at the root and the respective mutations along the edges of the associated tree. Note, that each edge in a tree might thus be annotated by multiple mutations, as it describes the evolutionary history of multiple genomic sites.

### 3.1 Implications of Our Design

Nodes in a genealogical forest represent unique subtrees (Section 3). Thus, queries using the trees’ topologies or the genetic sequences may easily reuse intermediate results on subtrees shared by *all* trees (*memoization*). In contrast, tree sequences allow for reusing intermediate results only among *adjacent* trees along the genome (see Section 1.5 for an explanation). We implement this memoization by storing the intermediate results in a lookup table, using the ID of the respective DAG node as the key. This memoization speeds up post-order traversals on the DAG, which we use to compute the LCAs (Section 3.3) and allele frequencies (Section 3.4), from which we derive numerous statistics in population genetics (Section 1.3).

### 3.2 Computing the Number of Selected Samples in a Subtree

For each query to the genealogical forest data structure, the user selects a subset of the forest’s samples to be considered (Section 1.3). Counting the number of selected samples which are part of each subtree of the genealogical forest (represented by a node in DAG) is an important building block for computing allele frequencies and the LCA. We define a post-order traversal on a DAG as iterating over all of its nodes such that all children of a node are processed before the node itself, thereby allowing us to reuse the intermediate result for these children using memoization (Section 3.1). Thus, to count the number of selected samples in each subtree, we assign *v*(*t*) = 1 for all tips *t* representing samples selected in the user query and *v*(*t*) = 0 to all other tips. Next, we perform a post-order traversal on the DAG by assigning to each node *n*_*k*_ the sum over the values assigned to its children *c*(*n*_*k*_), that is 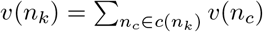 (Figure 2.b).

**Figure 2.**
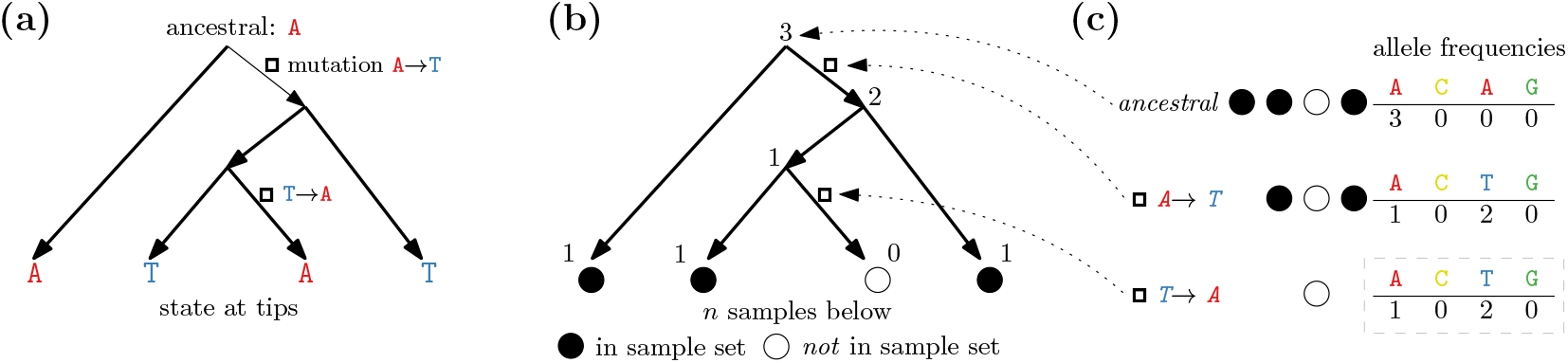
**(a)** Encoding a set of four sequences (here: one site) using a tree, an ancestral state, and mutations along the edges. **(b)** We compute the number of samples in the input sample set below each node using a post-order traversal. We need this information in order to **(c)** compute the allele frequencies (number of samples per genomic state) of each site by iterating over its mutations.

### 3.3 Computing the Lowest Common Ancestor

We compute the Lowest Common Ancestor (LCA, Section 1.2) of two or more tips in all trees represented by a genealogical forest using a variation of the algorithm described in Section 3.2. For this, we exploit that in a genealogical forest DAG, there exists exactly one path from each tip to each root. First, we assign each tip *t* selected as part of the query the tuple *n*(*t*) = (1, ∅) and all tips not selected the tuple *n*(*t*) = (0, ∅). Here, the first element of the tuple counts the number of selected samples which are part of the subtree represented by the node. We accumulate this counter up the trees (bottom up from the tips) using a post-order traversal on the DAG. Additionally, if for any node the sample counter is equal to the number of selected samples for the first time, this node must be the LCA for this subtree. Thus, we store the ID of this node in the second element of the tuple and propagate this value up the tree, instead of the sample count. Note, that we might select multiple nodes in the DAG as LCAs in a single query. However, as noted above, there is exactly one path from each tip to each root. This ensures that for each root in the DAG (representing a genealogical tree), we pick *exactly one* node that is reachable from this root as the LCA.

The runtime of this algorithm depends on the number of edges in the DAG, but not on the number of samples in the input sample set. Thus, computing the LCA of a large subset of the samples takes no longer than computing the LCA of only two samples (Section 6.2).

Note, that we could modify this algorithm to compute an almost-LCA-the lowest node under which “almost all” samples are located. This could increase the robustness of the biological interpretation against single “rogue” samples which were erroneously included in the input sample set and cause the LCA to be much higher than it would be without them.

### 3.4 Computing the Allele Frequencies and Derived Statistics

Most population genetics statistics depend on the allele frequencies [44], that is, how many `A`s, `C`s, `T`s, and `G`s we observe at a given site. We are interested only in variant sites, where at least two of these counts are *>* 0. Also, the allele frequencies at different genomic sites are evidently independent of one another. Again, we allow for selecting a subset of samples for a query. Given a query, we first count the number of selected samples contained in each subtree represented by the nodes in the genealogical forest DAG (Section 3.2). Let *S* = *{*`A`, `C`, `T`, `G`*}* be the set of alleles (here: possible genomic states) and *n*(*∗*) be the number of selected samples. For each site, we intend to compute the number of selected samples *n*(*x*) which exhibit allele *x ∈ S*. First, we set all samples to be in the ancestral state, that is *n*(*x*_ancestral_) = *n*(*∗*) and *n*(*x*) = 0 for all other *x* (Figure 2.c). Next, we iterate over the mutations at this site: Each mutation is associated with a subtree in the DAG and changes the state *x*_*i*_ ∈ *S* of all samples contained in this subtree to the state *x*_*j*_ ∈ *S*. We thus successively decrement *n*(*x*_*i*_) and increment *n*(*x*_*j*_) by the number of selected samples contained in the respective subtree.

From the allele frequencies, we now derive numerous population genetics statistics. The sequence *diversity* [39], for instance, reflects the probability that two random sequences differ at a random site (both chosen uniformly at random). *Segregating Sites* are sites exhibiting more than one allele (i.e., polyallelic sites). The *Allele Frequency Spectrum (AFS)* [17] is a histogram over the allele frequencies. Some statistics also work on multiple sample sets. For example the sequence *divergence* [39] is the probability that two sequences – each chosen uniformly at random from its own sample set – differ at any given site. More elaborate statistics can be derived from the above basic statistics, using up to four disjoint sample sets. Examples include *Patterson’s f*_2,3,4_ [42], *Tajima’s D* [52], and the *Fixation Index (F*_ST_) [58].

### 3.5 Memoizing on Shared Bipartitions

In contrast to computing the LCA, computing the allele frequencies is independent of the actual tree topologies as it requires only the sample bipartitions induced by the subtrees associated with the mutations. Remember, that these bipartition separate all tips of a tree into two disjoint sets (Section 2). For example, two identical mutations in two distinct subtrees ((*G*_0_,*G*_1_),*G*_2_) and (*G*_0_,(*G*_1_,*G*_2_)) will identically affect the respective allele frequencies. Memoizing on shared *bipartitions* allows reusing (slightly) more intermediate results than memoizing on shared *subtrees* (Section 6.5). Next, we detail the construction of the regular (unique subtrees) genealogical forest and this variant (unique bipartitions).

### 3.6 Constructing the Genealogical Forest DAG

In order to construct the genealogical forest DAG, we assign unique IDs to all subtrees of the genealogical trees forming the input. For this, let *h* be a pseudorandom hash-function. We start by assigning unique subtree IDs to the tips *T* = *{t*_0_, *t*_1_, …, *t*_|*T* |*−*1_*}* of the tree: id(*t*_0_) = *h*(0), id(*t*_1_) = *h*(1), …, id(*t*_|*T* |*−*1_) = *h*(*t*_|*T* |*−*1_). We then compute the unique subtree ID of each non-tip node by applying the bitwise exclusive or (⊕) over all of its children, followed by hashing the result using *h* (Figure 3.a). That is, for a node *n* with children *c*(*n*), we compute id(*n*) = *h*(⊕_*j∈c*(*n*)_id(*j*)). By using a symmetric function – i.e., id(*j*) ⊕ id(*i*) = id(*i*) ⊕ id(*j*) – we ensure, that the order by which we process the children does not affect the resulting subtree ID. In order for the actual topology of the subtree to influence its ID, we break the linearity of the ⊕ operator using a hash function. Otherwise, the ID would solely be determined by the bipartition induced by the respective subtree. For example, the following two subtrees would have the same subtree ID: `((A,B)C) & (A(B,C))`.

**Figure 3.**
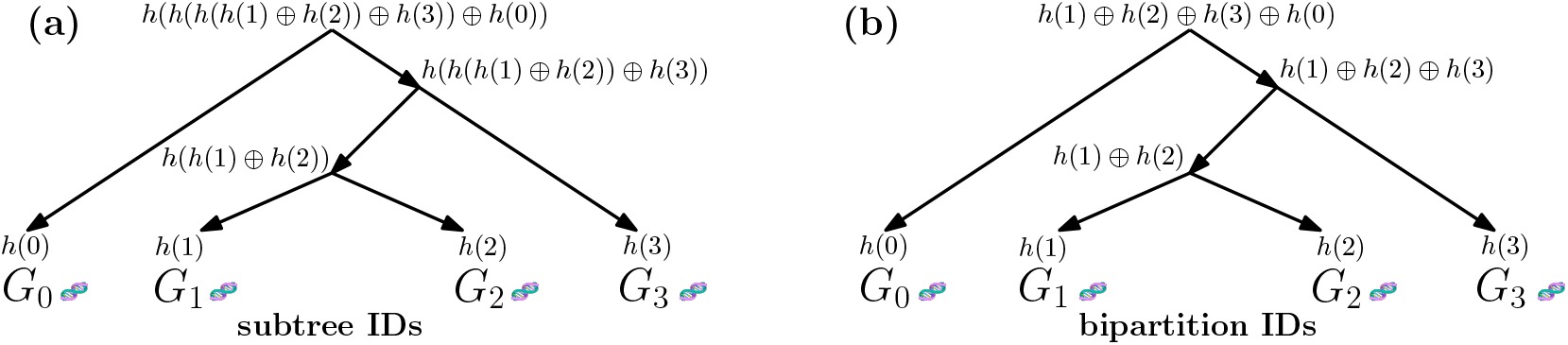
**(a)** Four sample genomes mapped to *t*_*i*_ *∈* 0, 1, 2, 3. The unique subtree IDs of the tips (i.e., samples) are *h*(*t*_*i*_) where *h* is a pseudorandom hash function. We compute the unique subtree IDs for non-tip nodes by applying *h* to the exclusive or (⊕) of all of its children. The resulting ID uniquely identifies a *subtree* including its topology. **(b)** Not breaking the linearity of the ⊕ results in a unique ID per set of samples in a subtree (i.e., not including its topology; called *bipartition*).

Using a 128 bit hash function, ensures a collision probability of 10*×*10^*−*16^ when computing *≈*3.8 *×* 10^11^ hashes [7]. This corresponds approximately to the probability of a single bit-flip occurring at any given second in the 128 bit needed to output a single subtree ID [49].

As noted in Section 3.5, the allele frequencies are invariant w.r.t. the actual topology of a subtree, and solely by the samples contained in it. For computing a unique *bipartition* ID, we therefore remove the linearity-breaking hash function *h* from the computation of the non-tip IDs, i.e., id(*n*_*i*_) = ⊕_*j∈c*(*n*)_id(*j*) for any non-tip node *n*.

#### Construction Algorithm

We construct the genealogical forest DAG from a given set of trees, by iterating over the input trees^8^ and processing the nodes of each tree in post-order. For each tree node (representing the subtree induced by it and its descendants) encountered during this post-order traversal, we compute the unique subtree ID and add the subtree to the DAG if it is not already present. For the sake of simplicity, we ensure that there exists exactly one distinct root in the DAG for each tree in the collection of input trees. Even if two trees are topologically exactly identical, each is represented by a separate root node in the DAG. They of course still share all subtrees below their respective, distinct, root node.

Recent versions of `tskit` provide information on which edges were inserted or removed when moving from one tree to the next along the genome. We exploit this information in order to recompute only those subtree IDs that might have changed, that is, all subtrees induced by nodes which are on the path from a changed edge to the tree root. Using this information results in speedups of 2 to 5, depending on the dataset.

Analogously to `tskit`, we encode the genomic sequence by storing the ancestral state as well as the respective mutations annotated at *nodes* of the DAG for each genomic site.

### 3.7 Balanced-Parenthesis Encoding of a Forest

Instead of encoding a genealogical forest as a DAG using explicit nodes and edges, an encoding extending the balanced parenthesis encoding for trees [27] with back-references is possible. This encoding is space-efficient and a post-order traversal over a balanced parenthesis encoded tree is a simple linear scan. Exploring the full design space of this approach and providing an efficient implementation represents an interesting challenge. However, initial experiments did not yield substantial speedups, and hence we do not repeat them here.

## 4 Experimental Setup

We implement our algorithms in C++20 and build our tool using `CMake` 3.25.1, `gcc` 12.1, and `ld` 2.38. We run our experiments on an AMD EPYC 7551P processor running at 2 GHz with 64 MiB of shared L3, 512 KiB of core-local L2, 32 KiB of core-local L1 data, and 64 KiB core-local L1 instruction cache. We use 8 banks of 32 GiB DDR4 RAM running at 2667 MT s^*−*1^. As our experiments are single threaded, only a single socket is being used. We compare `tskit` git rev `77faade5` and `gfkit` version `fbd2740`.^9^

## 5 Datasets

We evaluate our method on three freely-available empirical human (GRCh38) tree sequence collections (see Appendix for details): Thousand Genomes Project (TGP) [5] phase 3 autosomes, Simons Genome Diversity Project (SGDP) [35] autosomes, and the collection inferred by Wohns et al. [56] (“Unified”). We use all 22 autosomes for each of these collections in our experiments. We choose these specific tree sequence collections because to the best of our knowledge they constitute the only publicly-available empirical collections at present.

Additionally, we simulate a human dataset containing 640 000 samples (see Appendix for details). As we do not observe substantial runtime differences between distinct chromosomes of the empirical genomic data collections, we limit our simulated data benchmarks to chromosome 20 to conserve computation resources and reduce our CO2 footprint.^10^

## 6 Evaluation

We evaluate the implementation of our proposed data structure “genealogical forests” (`gfkit`) regarding query speed (Section 6.1 and Section 6.2) and storage space used (Section 6.6). Additionally, we assess the algorithmic reasons for the obtained speedups (Section 6.4 and Section 6.3) and report the time required to convert `tskit` tree sequences into the `gfkit` data structure (Section 6.7).

### 6.1 Speedup for Computing Statistics Based on the Allele Frequencies

We evaluate the runtime of our proposed genealogical forest data structure and its associated implementation `gfkit` by comparing it to the state-of-the-art reference implementation of tree sequences, `tskit`. We benchmark the runtimes of various important statistics in population genetics (Section 1.3), including statistics that are based on the allele frequencies (e.g., AFS) as well as on the topology (LCA). We use 10 repeats for each runtime measurement on each of the 22 autosomal chromosomes of each empirical collection (TGP, SGDP, and Unified; see Section 5) and on chromosome 20 of the simulated dataset. The mean standard deviation of runtimes across the 10 repeats of each statistic, collection, and chromosome is below 1.3 %. We report a median speedup of 4.0 of `gfkit` (ours) over `tskit` (state-of-the-art) for computing various allele frequency-based statistics (Figure 4). The absolute runtimes range from 775 to 1770 ms for `tskit` and 152 to 340 ms for `gfkit`.

**Figure 4.**
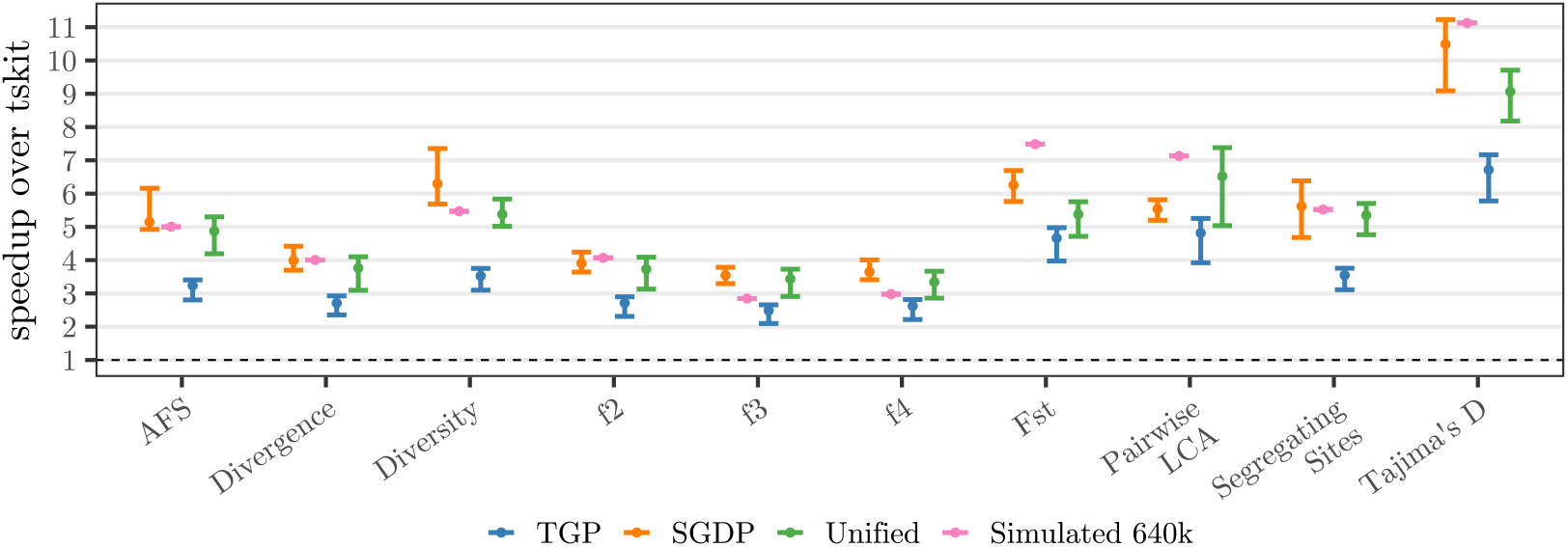
Speedup of `gfkit` (ours) over `tskit` (state-of-the-art) for computing statistics on three empirical and one simulated dataset (human). The bars indicate the speedup range and the dot the median speedup across all chromosomes of the respective collection. We use all 22 autosomal chromosomes from the Thousand Genome Project (TGP), Simons Genome Diversity Project (SGDP), and the Unified collection. We use chromosome 20 of a simulated human collection containing 640 000 samples. AFS: Allele Frequency Spectrum. f{2,3,4}: Patterson’s *f*. Fst: Fixation Index.

Profiling using Intel VTune shows that `gfkit` spends over 90 % of its runtime in the post-order traversal during the computation the allele frequencies. The numerical computations for the actual statistics thus amount for less than 10 % of runtime. Therefore, the number of nodes in the genealogical forest DAG, as well as the in-degree of those nodes appear to constitute the dominating runtime factors. These values correspond to the number of unique subtrees in the genealogical trees used to construct the genealogical forest, and the number of times that an intermediate result can be reused during the post-order traversal, respectively. Thus, we provide more details on these measurements (Section 6.3 and Section 6.4).

### 6.2 Speedup for Computing the Lowest Common Ancestor

We compare `gfkit`’s and `tskit`’s runtimes for computing the Lowest Common Ancestor (LCA) of two (“pairwise”) or more samples. The asymptotic runtime of `gfkit`’s LCA-algorithm (Section 3.3) does not depend on the number of selected samples. In contrast, given three or more samples, `tskit` performs an analogous number of LCA-queries: First, it computes the LCA of two arbitrary samples, which it subsequently uses as input for the next LCA query; together with another sample from the input. After processing all samples, `tskit` obtains the overall LCA. Therefore, `tskit`’s LCA-algorithm scales linearly with the number of samples in the input.

We report the speedups of `gfkit` over `tskit` when computing the LCA (Figure 4 and Figure 5). In order to save computational resources and reduce our environmental footprint, we perform only 3 repeats when computing the LCA of more than two samples using `tskit` (runtime up to 34 min). We observe speedups of `gfkit` over `tskit` of 5.5 (median) for pairwise queries, 208 (median) when selecting 10 % of the samples, and 990 (median) when selecting 50 % of the samples as input sample set. Experiments on the simulated dataset took 64 min for `gfkit` (median 155 ms per query) but did not finish for `tskit` in a week.

**Figure 5.**
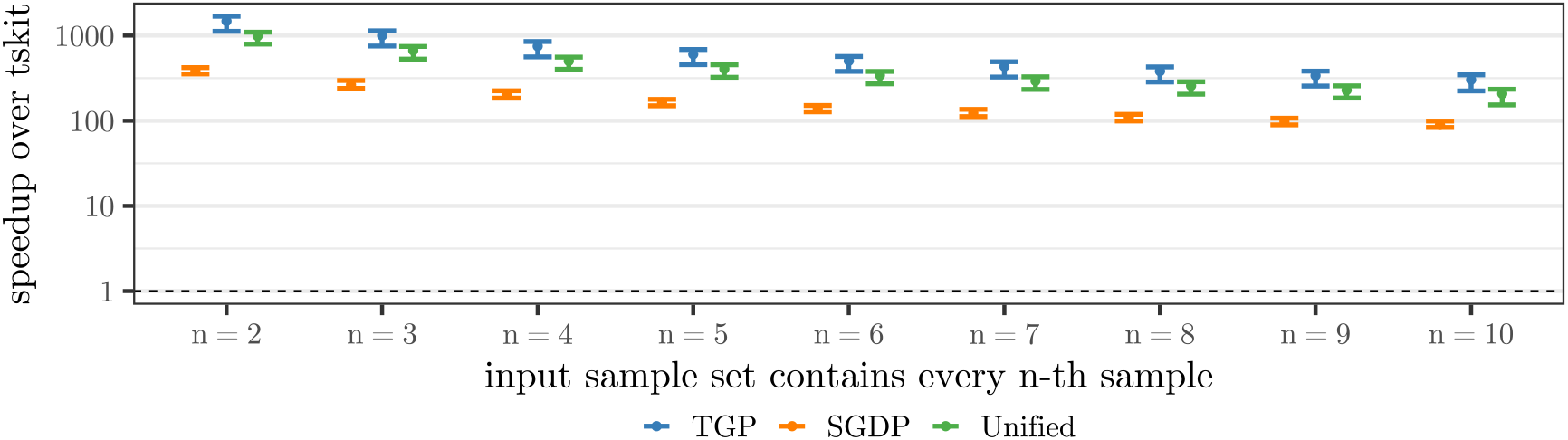
Speedup of `gfkit` (ours) over `tskit` (state-of-the-art) for computing the Lowest Common Ancestor (LCA) of every *n*-th sample in the dataset. The runtime of `gfkit`’s LCA-algorithm does not depend on the number of samples in the sample set. In contrast, the runtime of `tskit`’s LCA-algorithm depends linearly on the number samples in the sample set.

### 6.3 Proportion of Subtrees that are Unique

Each query spends the majority of its time in the post-order traversal. Thus, the number of nodes and edges in the `gfkit` DAG is *the* determining runtime factor. The number of unique subtrees in the input tree set determines the number of nodes in the DAG. We report the number of overall subtrees in the input as well as the absolute and relative number of unique subtrees (Table 1). For example, in the TGP collection, 0.48 % (mean) of the subtrees are unique per chromosome. Additionally, we report that the absolute number of unique subtrees in a simulated dataset (human, chromosome 20) with 640 000 samples is not substantially higher than for the shown empirical collections containing 554 to 7508 samples, respectively.

**Table 1.** Number of overall and unique subtrees and the proportion of subtrees that are unique. The ranges given cover all 22 autosomal chromosomes of the respective collection. We report the arithmetic mean and standard deviation across all chromosomes of a collection.

| Collection | Chr. | Samples | Subtrees<br>$\times 10^6$ | Uniq. Subtrees<br>$\times 10^6$ | Uniq. Subtrees /<br>Subtrees |
| --- | --- | --- | --- | --- | --- |
| SGDP | all | 554 | 84 to 587 | 1.8 to 11 | $0.020 \pm 0.001$ |
| TGP | all | 5008 | 2150 to 14 916 | 11 to 69 | $0.0048 \pm 0.0003$ |
| Unified | all | 7508 | 698 to 2708 | 3.2 to 18 | $0.006 \pm 0.002$ |
| Sim. 640k | 20 | 640 000 | 695 185 | 14 | 0.000 02 |

In conclusion, only 0.002 to 2 % of the subtrees in the tested collections are unique. This proportion decreases as we add more samples (and thus subtrees) from the same species. This is due to the absolute number of unique subtrees not increasing substantially for larger collections, thus, neither does the runtime of the queries. The number of mutations per subtree is 3 to 4 orders of magnitude smaller^11^ for the 640 000-sample dataset compared to the collections with *≤* 7508 samples. Therefore, there is less signal to resolve the evolutionary history of the samples, possibly leading to larger unresolved subtrees. It remains an open question how other data characteristics (for example the species when not considering human populations) influence the number of unique subtrees.

### 6.4 Reusing Shared Subtrees

Apart from to the number of nodes in the `gfkit` DAG, the number of edges also influences the runtime of the post-order traversal. However, the proportion of edges to nodes also serves as an indicator for memoization performance. During the post-order traversal, the in-degree of a node is equal to the number of times its result is used (and reused). We report the mean in-degree across all non-root nodes in all trees across all chromosomes of each collection (Appendix). Thus, each intermediate result on a unique subtree is reused on average 4.12 (SGDP), 5.48 (TGP), 5.59 (Unified), or 5.10 (Simulated 640k) times.

### 6.5 Speedups when Memoizing on Shared Bipartitions

We also benchmark queries on the bipartition-DAG described in Section 3.5. Here, we observe a median speedup of 4.7 over `tskit` across all empirical collections (Appendix). Queries on the bipartition DAG are on average 1.14 *±* 0.09 (mean *±* sd) times faster than on the subtree-DAG. There are 5 to 20 % fewer unique bipartitions than there are unique subtrees; corresponding to fewer nodes in the bipartition-DAG. The average in-degree in the bipartition-DAG, and thus number of times we reuse an intermediate result, ranges from 4.44 to 5.20 (Appendix), compared to 4.12 to 5.49 on the subtree-DAG. These two measurements explain the (moderately) faster runtimes of the bipartition-DAG compared to the subtree-DAG. However, as the bipartition-DAG does not support topology-aware queries (e.g, LCA), we do not consider the tradeoff worthwhile with respect to the increased code complexity and storage unless additional optimizations further reduce the runtimes.

### 6.6 Storage Space Needed for Encoding the Forest

In a sense, the genealogical forest DAG factors-out all unique subtrees that a tree sequence describes using edge insertions and removals. Concerning the space usage, there are two contrary effects at play here: On the one hand, tree sequences store some identical subtrees multiple times, but using distinct edges and/or nodes. On the other hand, a single edge insertion or removal can induce multiple new subtrees along the path from the insertion/removal point up to the tree root. In the first case, tree sequences are the more space-efficient representation, in the second case, genealogical forest are.

In order to quantify the trade-off between the two effects, we compare the size of the genealogical forest DAG against the size of the respective tree sequence. However, the tree sequence implementation `tskit` stores a variety of additional meta-data, to support operations which we currently do not implement in `gfkit`. Additionally, `tskit` does not include some optimizations, for example, not explicitly storing node or edge IDs. We thus perform a theoretical, instead of an empirical space requirement comparison.

A tree sequence as well as a genealogical forest could minimally be described by its (directed) edges plus sequence information. Let *ι* be the number bits needed to encode a node ID and *ϕ* be the number of bits needed to encode a position in the genome. Thus, choosing an edge list (see below) we need 2 · *ι* bit for each edge in a genealogical forest. In tree sequences, an edge is valid for a specified region of the genome, requiring 2 *· ϕ* additional bits per edge to store this information. Note, however, that the number of nodes and edges differs between `tskit` and `gfkit` and these numbers are thus not directly comparable. Additionally, we cannot arbitrarily sort `tskit`’s edges without incurring a performance penalty, as we require the insertion order in order to efficiently move from one tree to the next along the genome.^12^ In `gfkit`, all outgoing edges of a node could be stored consecutively in an edge list, which would allow omitting some of the `from` node IDs, conceptually creating an adjacency array.

Both, `tskit` and `gfkit`, describe the genomic sequences using one ancestral state and a list of mutations per genome site. We can encode each mutation using a site ID, the node ID at which the mutation occurs, plus the derived state. Let *s* be the number of sites, *σ* be the number of bits per site ID, *γ* be the number of bits per genomic state, and *m* be the overall number of mutations. The sequence information thus (theoretically) requires *s · γ* + *m ·* (*σ* + *ϕ* + *γ*) bit. In practice, `tskit` and `gfkit` store additional information: For example, in order to avoid traversing up the tree to the parent mutation (or root node), `tskit` stores a pointer to the parent mutation (64 bit) and `gfkit` stores the parent mutation’s state (2 bit). However, we omit these and other implementation-specific constants here.

For our calculations (Table 2), we assume *ι* = *ϕ* = *σ* = 32 bit integers for node IDs, site IDs, and genomic positions^13^ and four possible genomic states, thus *γ* = 2 bit. We conclude that the edge removals and insertions used to store the set of genealogical trees by `tskit` are more-space efficient than `gfkit`’s method of storing a node for each unique subtree. However, we argue that, as the differences are in the same order of magnitude than implementation-specific constants (e.g., choice of data-types or explicitly storing IDs), one could mitigate against this effect. Therefore, it remains a tradeoff between query-speed and space usage. It is subject of future work to evaluate the genealogical forest variants using the compressed edge list (above) and the balanced parenthesis representation (Section 3.7).

**Table 2.**
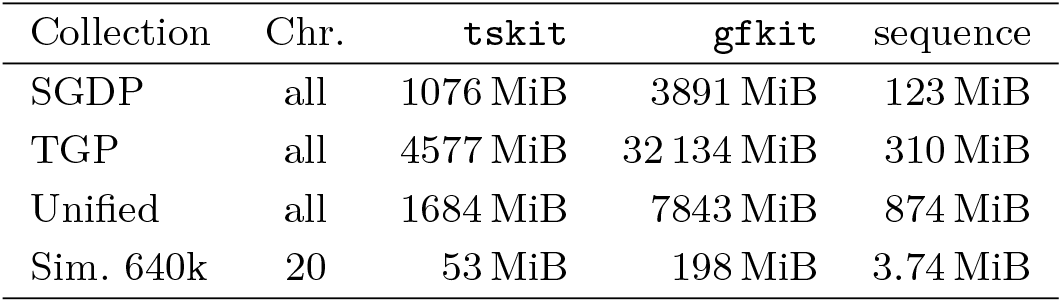
Theoretical space usage of `tskit` vs `gfkit`. The description of the trees is substantially larger than the description of the sequence (ancestral states and mutations). `tskit` encodes the trees as a tree sequence (Section 1.5), whereas `gfkit` encodes them using a genealogical forest (Section 3).

### 6.7 Converting Tree Sequences to Genealogical Forests

To the best of our knowledge, there is no fundamental reason why tools could not output genealogical forests instead of tree sequences. Currently, however, these tools output the established `tskit` format – and will probably continue doing so for a while, since the current implementation of our data structure, does not support all features of `tskit` yet. While we provide corresponding `gfkit` files (Appendix) for the openly available `tskit` collections (Section 5), the time taken to convert `tskit` to `gfkit` input files is not negligible in general.

Kelleher et al. [30] report tree sequence inference times of 5 min for chromosome 20 of SGDP (Section 5) and 2 h for chromosome 20 of TGP using 40 cores. We convert their output files from the `tskit` to the `gfkit` format on a single core in 6.5 s and 123 s respectively. We thus argue that one can convert empirical collections in a negligible amount of time compared to the time required for inferring them. Additionally, in principle, tree sequence inference could directly output the `gfkit` format, thereby entirely eliminating the conversion step.

## 7 Numerical Stability

Basic operations such as additions or multiplications on IEEE 754 floating point numbers always induce rounding errors [19]. The magnitude of these errors directly depends on the difference order of magnitude between the two operands. Thus, computing running sums, where we sum many small values to an ever-growing sum, is particular rounding-error prone.

We often compute the statistics in population genetic on a per site (Section 1.3) basis and then sum over the values for all sites to compute an average. Doing this using a running sum – as it is currently implemented in `tskit` and `gfkit` – leads to increasing round-off errors the more variant genetic sites there are. Other statistics, for example, Patterson’s *f*_4_ entail the multiplication of the number of samples in four different sample sets. On datasets with more than 262 143 samples^14^, the result of this product might be too large to be represented in as a 64 bit integer. We (for the sake of result verification), as well as `tskit`, thus choose to represent these products as 64 bit IEEE 754 floating point numbers, thus introducing potentially substantial numerical errors.

It is currently unknown if errors introduced by these simplifications substantially influence the biological interpretation of the results. Further, rounding errors imply that the order of operations influences the result, even though the input operands are bit-identical, thus impacting reproducibility. Even when the software version and the random seeds are fixed, different algorithms, compiler optimizations, or degrees of parallelism influence the order of operations. Reports on these effects influencing the results of scientific computations exist for example in the fields of phylogenetics [13], sheet metal forming [15], and fluid simulations [54].

Therefore, a thorough analysis of the potential impact of induced numerical errors on standard population genetics statistics with respect to their effect on downstream analyses as well as on reproducibility of results is required. Ideally, tool authors would agree upon a standardized order of calculations, such that all software in the field yields comparable and reproducible results with well-understood numerical inaccuracies. Further, the results obtained should be invariant under addition of further – but ignored in the query at hand – genomes and mutations to the dataset. As the numerical calculations require less than 10 % of the overall runtime, we are optimistic that even introducing arbitrary precision calculations would result in only a negligible runtime overhead.

## 8 Conclusion and Future Work

In recombining organisms, a set of genealogical trees is often used to describe the evolutionary history of the studied samples. While tree sequences (`tskit`) compress these trees using edit operations from one tree to the next along the genome, genealogical forest (`gfkit`) compresses them into a DAG where each node represents a unique subtree of the input (Section 3). While the genealogical forest encoding (theoretically) requires a factor 3.6 to 7.0 more space than tree sequences (Section 6.6), it also yields speedups by a factor of 2.1 to 11.2 (median 4.0) when computing pairwise LCA queries and important statistics in population genetics (e.g., sequence diversity). In contrast to `tskit`’s LCA-algorithm, the runtime of `gfkit`’s LCA-algorithm does not depend on the number of samples selected in the query. Thus, `gfkit`’s LCA queries are substantially faster, e.g. a speedup of 208 (median) when querying 10 % of samples and a speedup of 990 (median) when querying 50 % of samples. Additionally, we describe, implement, and benchmark an alternative data structure, which represents the set of genealogical trees as a DAG in which each node represents an unique bipartition in the input. This variant is slightly faster but does not support topology-aware queries (e.g., LCA). To explain this, we show, that subtrees containing the same samples but having a different topology are actually rare in the analyzed datasets.

As more genomes are being sequenced, future datasets will contain a multiple of today’s samples. However, we observe speedups (and runtimes) comparable to the largest existing empirical datasets for a simulated human dataset with 640 000 samples (Section 6.1). We show, that this might be due to the number of unique subtrees being comparable to those of current empirical datasets with at most 7508 samples, thus resulting in DAGs of about the same size. As the post-order traversal on the genealogical forest DAG takes up over 90 % of a query’s runtime, the size of this DAG is the determining runtime factor. Which characteristics of the input influence the number of unique subtrees remains an open question.

`Tskit` was not build in a day and neither was `gfkit`. Currently, `gfkit` does not support all `tskit` features, yet most of these less common features should be straightforward to implement. For example, `gfkit` currently supports only sample weights of 0 and 1 (a sample is in the sample set, or it is not), as these are sufficient for implementing the allele frequencies and derived statistics as well as LCA queries. Further, `gfkit` currently supports only site statistics on the entire genome, that is, neither branch- nor node-based statistics as well as no sliding windows over the genomic sites. Additionally, `gfkit` assumes that for each genomic site, a single tree describes the evolutionary history of all tips, while `tskit` allows for multiple (partial) trees.

Next to lifting these limitations, additional improvements include evaluating a top-tree based genealogical forest (Section 2) and extending LCA queries with more than two samples to also report ancestors which are common to “almost all” selected samples, thus making the query robust against single samples erroneously included in the query (Section 3.3).

Tree sequences opened up new possibilities of storing and processing large sets of genealogical trees and sequences used in population genetics. Genealogical forests provide a substantial reduction in runtime and a possibly more intuitive representation over the state-of-the-art. We believe that these improvements will boost the development of new analyses in the field of population genetics, for example they might be used for developing appropriate optimization criteria and techniques for automatically determining subpopulations.

## 9 Acknowledgments

This project received funding from (a) the European Research Council (ERC) under the European Union’s Horizon 2020 research and innovation program (grant agreement No. 882500) (b) the European Union (EU) under Grant Agreement No 101087081 (Comp-Bio-div-GR). (c) the Ministry of Science, Research, and the Arts of Baden-Württemberg (Az: 33-7533.-9-10/20/2) to Peter Sanders and Alexandros Stamatakis.

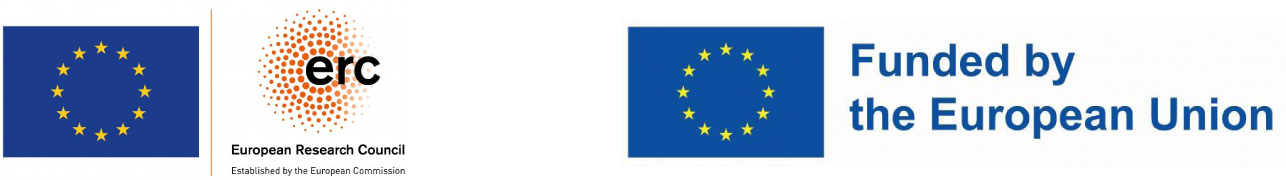

## A Dataset Details

Tree sequence datasets were inferred using `tsinfer` [30] version `0.2.1` and dated using `tsdate` [56] version `0.1.4`. The description on Zenodo is incorrect; see: https://tskit-dev.slack.com/archives/C010834D669/p1713340040404519. We simulate the “Sim. 640k” dataset containing 640 000 samples using `stdpopsim 0.2.0`^15^ and the `HapMapII_GRCh38` genetic map. We convert tree sequence to genealogical forest files via `gfkit` version `fbd2740`.

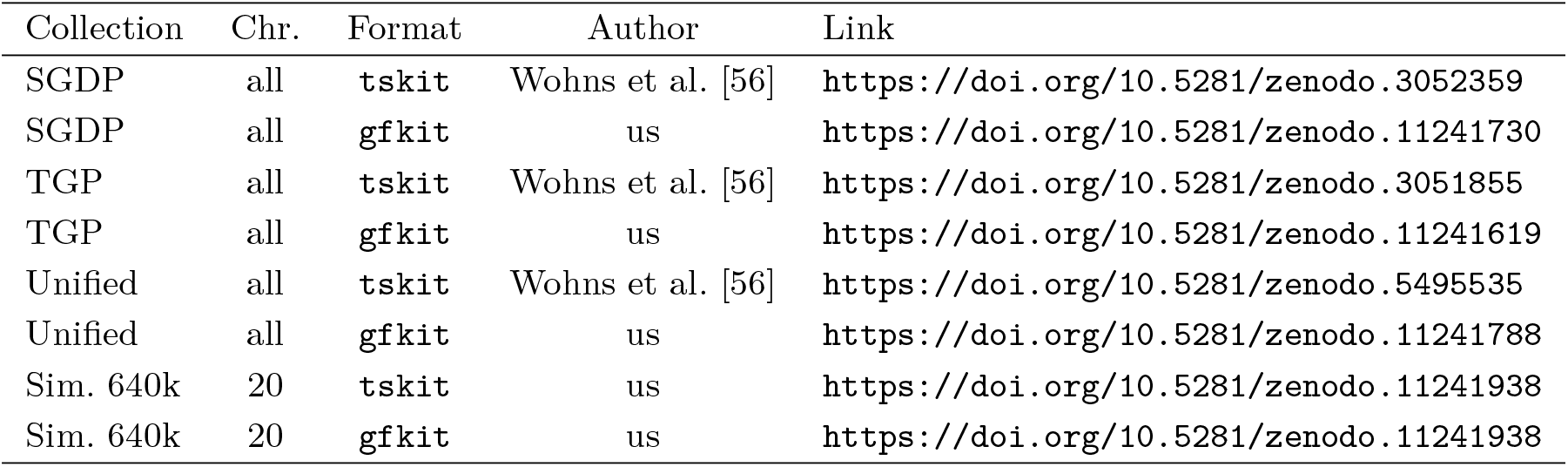

## B Reusing Shared Bipartitions

**Table 3.** Average in-degree of non-root nodes in the `gfkit` DAG. The in-degree of a node in the DAG is equal to the number of times its intermediate result is reused during the post-order traversal.

| Dataset | Chr. | Mean In-degree<br>Bipartition | Mean In-degree<br>Subtree |
| --- | --- | --- | --- |
| SGDP | all | 4.44 | 4.12 |
| TGP | all | 5.89 | 5.48 |
| Unified | all | 5.20 | 5.49 |
| Sim. 640k | 20 | — | 5.10 |

<sup>15</sup>https://github.com/popsim-consortium/stdpopsim

## C Speedups Using the Bipartition-DAG

**Figure 6.**
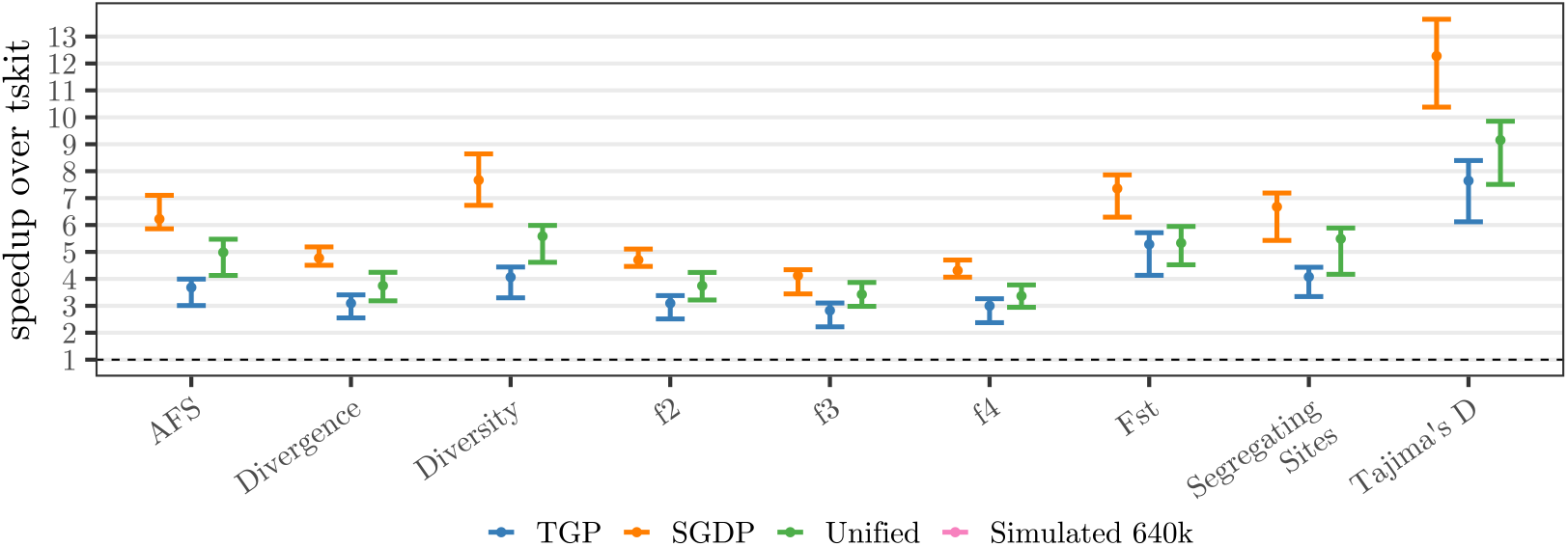
Speedup of `gfkit` (ours) over `tskit` (state-of-the-art) when computing various statistics on three empirical and one simulated dataset (human). The bars indicate the range of speedups and the dot the median speedup across all chromosomes of the respective collection. We use all 22 autosomal chromosomes for Thousand Genome Project (TGP), Simons Genome Diversity Project (SGDP), and the Unified collection. We use chromosome 20 of a simulated human dataset with 640 000 samples.

## Footnotes

1 In this work, human always refers to *homo sapiens*.

2 Also called the Most Recent Common Ancestor (MRCA).

3 https://digital-strategy.ec.europa.eu/en/policies/1-million-genomes

4 For some problems, the minimum work required might not be known.

5 Minimal tree grammars are 𝒩𝒫-hard to construct and they do not support efficient navigation [9].

6 Branch lengths are irrelevant for genetic variation based statistics and the LCA, thus we omit them.

7 Invariant sites do not carry evolutionary information and are thus filtered out during pre-processing.

8 The order of trees in not relevant.

9 https://github.com/tskit-dev/tskit and https://github.com/lukashuebner/gfkit

10 Wohns et al. [56] use chromosome 20, as they consider it representative for genome-wide patterns.

11 Mutations per subtree: *≈* Sim. 640k: 0.000 000 7, SGDP: 0.002, TGP: 0.0002, and Unified: 0.003

12 We could determine the deletion order on the fly by using a priority queue containing only the currently active edges.

13 This is true for `gfkit`. However, `tskit` uses *ϕ* = 64 bit IEEE 754 floating points for genomic positions.

14 When the four sample sets have the same size: (262144*/*4)^4^ = 2^64^

15 https://github.com/popsim-consortium/stdpopsim

